# A piece of evidence for pathogenesis of cerebral small vessel disease caused by blood-brain barrier dysfunction —— Based on the STRIVE-2 imaging features

**DOI:** 10.1101/2025.02.24.640002

**Authors:** Yi Yang, Huide Ma, Bowei Liu, Rong Wu, Sujuan Sun, Xia Li, Yu He, Shumin Wang

## Abstract

**Background:** Blood-brain barrier (BBB) dysfunction caused by endothelial cell injury is one of the widely accepted pathogenesis of small cerebral vessel disease (CSVD). However, the early microcirculation studies were notably deficient in providing compelling imaging evidence, which has hampered a comprehensive understanding of the pathophysiological processes underlying BBB dysfunction in CSVD. The onset of CSVD is insidious and the clinical manifestations are diverse, diagnosis of CSVD relies primarily on neuroimaging information currently. These MRI imaging features mainly represent the end-stage brain parenchyma injury, rather than the vascular lesion itself. Therefore, clarifying the pathophysiological processes related to cerebral microcirculation in CSVD is of great significance for early treatment intervention of CSVD.

**Methods:** Twenty rats were used to prepare CSVD animal model by ultrasound combined with ultrasonic microbubble contrast agent. MRI, functional ultrasound (fUS) imaging and ultrasound localization microscopy (ULM) were used to evaluate microcirculation changes in acute and chronic models.

**Results:** Currently, our research has successfully validated that the imaging characteristics of this model align with the four established criteria outlined in the STRIVE-2 classification. FUS shows increased microflow velocity in the molding area, which can be used for initial evaluation of CSVD models in the acute phase. ULM has revealed a progressive reduction in the density of both the short cortical arteries and the long medullary arteries, along with their respective branches, over time. Concurrently, there has been a noted deviation in blood flow velocity.

**Conclusion:** This model offers compelling evidence at the microvascular level, suggesting that blood-brain barrier (BBB) dysfunction is a central pathophysiological mechanism in the etiology of CSVD. Then it provides a straightforward, efficient, and widely applicable tool and continuous monitoring method for the further investigation of CSVD from the macroscopic to the microscopic circulation level, with significant long-term implications for CSVD diagnosis and treatment.

## Introduction

With the development of medical imaging, people have a preliminary understanding of cerebral small vessel disease (CSVD) and its pathogenesis^1-3^, especially vascular endothelial dysfunction may be involved in the occurrence mechanism of CSVD^4,5^, and the destruction and activation ability of endothelial cell are the current research hotspots^6^. The onset of CSVD is insidious and the clinical manifestations are diverse, and it is difficult to obtain microvascular information in the acute stage^7,8^. Diagnosis of CSVD relies primarily on neuroimaging information^9^. In 2023, 50 international experts and 4 external consultants published an updated version (STRIVE-2) of the standards for reporting vascular changes on neuroimaging in Lancet Neurology^10^, which has become an internationally recognized standard for diagnostic imaging and research of CSVD. Imaging features of STRIVE-2 include recent small subcortical infarct, lacunes (of presumed vascular origin), white matter hyperintensity (of presumed vascular origin), perivascular space, cerebral microbleeds, cortical superficial siderosis, intracerebral hemorrhage brain atrophy and other hemorrhagic signs. These MRI imaging features mainly represent the end-stage brain parenchyma injury^11^, rather than the vascular lesion itself, and mainstream scholars believe that the microvascular injury precedes the brain parenchyma injury^12^. Therefore, clarifying the pathophysiological processes related to cerebral microcirculation in CSVD is of great significance for early treatment intervention of CSVD.

Blood-brain barrier (BBB) dysfunction caused by endothelial cell injury is one of the widely accepted pathogenesis of CSVD^4,5,13,14^. Based on this, we used ultrasound combined with ultrasound contrast agent to damage the microvascular endothelial cells in the mold area, causing BBB dysfunction, and simulating the pathogenesis of cerebral small vessels, so as to prepare a rat model of cerebral small vessel disease^15^. This model provides a new modeling method for the study of CSVD, and finds that ultrasound localization microscopy (ULM) is an effective evaluation method, which can be directly used to evaluate the hemodynamic changes of CSVD. Based on the mechanism of BBB dysfunction caused by endothelial cell injury in CSVD, this paper used various modalities such as MRI imaging features, ultrasound functional imaging (fUS)^16^ and ULM to continuously observe the microcirculation changes of early CSVD after modeling. Continuous observation of the microcirculatory changes in early CSVD after modeling provided imaging evidence at the microcirculatory level for the mechanism of CSVD formation due to dysfunction of BBB.

## Materials and Methods

### 1. Rat model of CSVD

This study was approved by the local ethics committee of Peking University Third Hospital (A2023122). 20 Adult male Sprague-Dawley rats (Beijing Vital River Laboratory Animal Technology Co., Ltd.). 40 bottles Sonoclar (Beijing Ferrymed Co., Ltd). For molding, please refer to our preliminary work for specific operation methods^15^. Ultrasound imaging and MRI were observed in the acute stage, and MRI and H&E staining were observed in the chronic stage.

### 2. Functional ultrasound

Functional ultrasound imaging acquisitions were performed with a functional ultrasound scanner (Iconeus One, Iconeus, Paris, France) equipped with a linear array transducer (15 MHz central frequency, 128 elements, 110 μm pitch). Composite frames (11 angles between -10° and 10°) were obtained at 500 Hz. A singular value decomposition filter (removing the first 60 singular values) was used to separate the blood signal from the tissue and summation to obtain a power Doppler image. Between the two images, a 0.2-second pause is added to allow the motor to move to the next slice. Twenty-one coronal planes were imaged every 0.2 mm among 4 mm (from β-3 mm to β+1 mm) at planar resolution of 110 μm × 100 μm. The data were saved with IconScan software and exported with IconStudio as a video file. Postprocessing was performed with MATLAB scripts, and a graphical user interface program was developed.

### 3. Ultrasound Localization Microscopy

ULM was also performed on the Iconeus One system. 100 μL Sonoclar microbubbles solution (diluted with saline by 1:9) was injected to the tail vein before image acquisition. The ultrafast compounding (7 angles between -6° and 6°) plane wave sequence was then performed with the frame rate of 1000 Hz. 90 seconds raw data were recorded for each acquisition. The superloc software provided by Iconeus were used to compute and display the ULM images, and the density and velocity maps were exported as image files with a resolution of 11 μm × 10 μm. Postprocessing of the maps was also performed with MATLAB scripts we developed.

### 4. Magnetic Resonance Imaging

Magnetic resonance imaging (MRI) was performed 24 hours and 3 weeks later. After anesthesia with pentobarbital sodium (300 mg/kg), the rats were placed in prone position in a clinical MRI, FLAIR T2-weighted images were acquired on the Discovery™ MR750 advances 3.0T scanner (GE Discovery MR 750, GE Healthcare, Milwaukee, WI, USA) using a brain coil, typical imaging parameters were: slice thickness = 1.5 mm (Gap = 1 mm), TR/TE = 10000/84.85 ms (FA = 111°), TI = 1658.5 ms, NSA = 1, reconstruction resolution = 0.16 * 0.16 mm2, image matrix = 256 * 256.

### 5. Histological diagnosis

After anesthesia perfusion, the rats were dissected and the coronary brain sections were prepared, fixed with 10% formalin buffer and embedded with paraffin. A rat brain section was stained with hematoxylin and eosin (H&E) for histopathological evaluation, and the section was observed by optical microscope.

## Results

### 1. Comparison of the CSVD model with STRIVE-2 imaging features

Fig. 1(a) is an MRI T2-weighted imaging of the coronal section of the rat brain 30 days after modeling, the solid arrow showed the cerebral cortex in the modeling area, which was thinner than the contralateral side, and the hollow arrow showed the paraventricular white matter area, which was hyperintense compared with the contralateral side. Fig. 1(a) is consistent with cerebral atrophy and white matter hyperintensity in STRIVE-2. Fig. 1(b) is an MRI T2-weighted imaging of the sagittal section of the rat brain one day after modeling, and the hollow arrows show the mixed area of high and low signals, and the peripheral of the mixed area can be seen with high signal wrapping. Fig. 1(c) was the ULM image one day after modeling, and the cerebral cortex blood vessels shown by the hollow arrow are less than those on the contralateral side, which can be consistent with the performance of brain microinfarction in STRIVE-2 when combined with Fig. 1(b). Fig. 1(d)-(e) are the brain H&E staining one day after modeling, a small bleeding area can be seen in the area shown by the hollow arrow in the Fig. 1(e), and the H&E staining in the adjacent area is faint, which can be consistent with the cerebral microhemorrhage in STRIVE-2 and the edema of the surrounding brain tissue after microhemorrhage.

**Figure 1.**
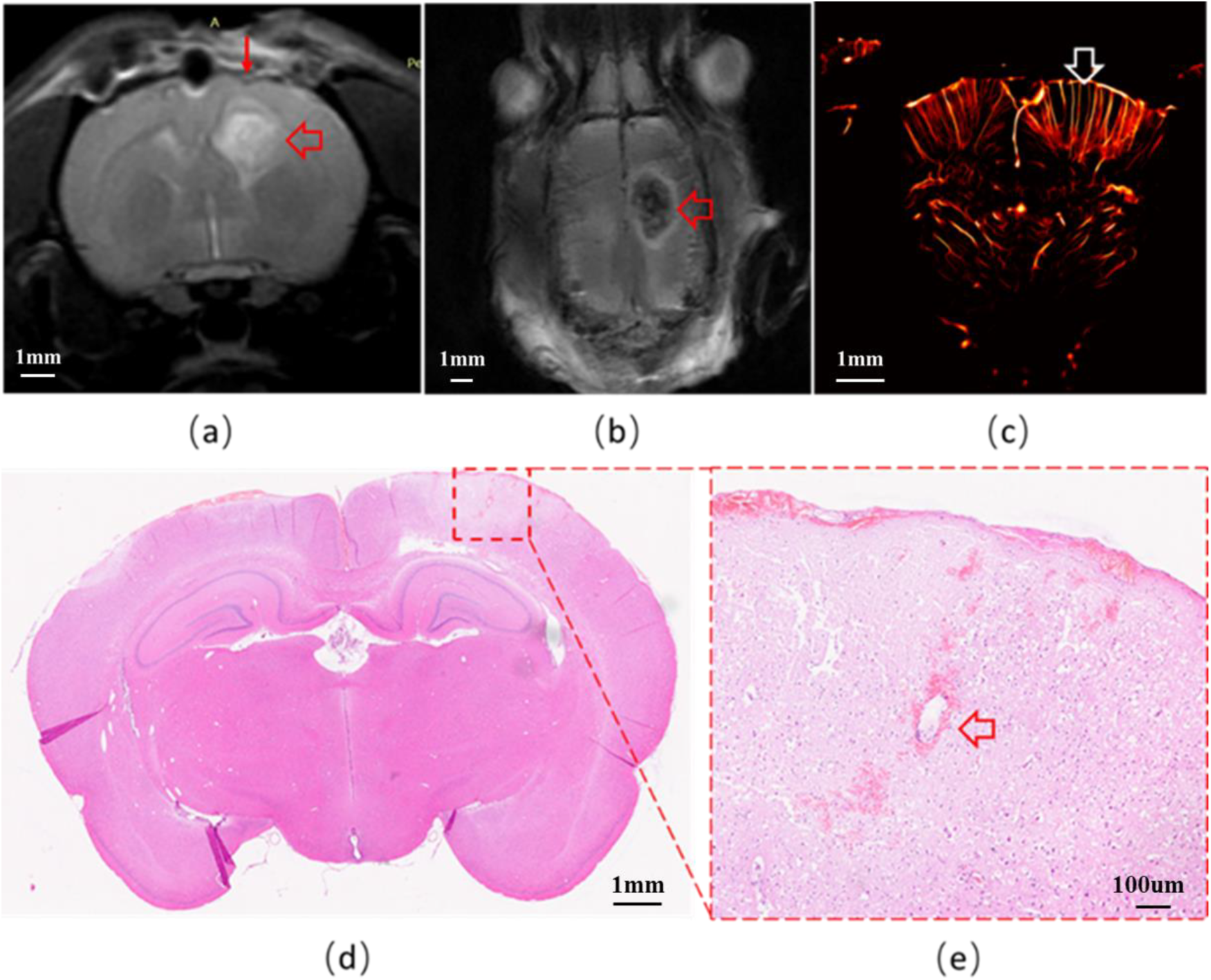
(a) T2-weighted MRI image with line arrows showing cortical region thinning and hollow arrows showing white matter hyperintensity. (b) T2-weighted MRI image with hollow arrows showing mixed areas of high and low signals. (c) Ultrasound positioning microscopic image, with sparse blood vessels in the irradiated area shown by the hollow arrow. (d) is the H&E staining plot, (e) is the magnification of the irradiated area, and the hollow arrow shows the microhemorrhage area.

### 2. Preliminary evaluation of the acute phase of fUS in the CSVD model

Fig. 2(a) is the fUS image of the coronal section of the rat brain one day after modeling, and the micro blood flow signal in the blue ellipse can be seen higher than that on the opposite side, indicating that the blood flow velocity in this area is increased, the molding region exhibits an ellipsoidal shape. Fig. 2(b) is the gray distribution of mean values of each vertical pixels along the white arrow direction in the yellow box of Fig. 2(a), and the mean gray values along the arrow direction is increased. As observed in Fig. 1(b) and Fig. 2(a), the modeled region approximates an ellipsoid, hence the variation of the cross-sectional area along the coronal direction follows a parabolic curve. Fig. 2(c) presents the quantitative assessment of the high-signal area in consecutive sections of the modeled region. By fitting a curve and calculating the area under the curve, we have determined the volume of the modeled region to be V=1.16 mm^3^. This method, with minimal craniotomy-related trauma, provides a volumetric quantitative approach for further research on the modeled region.

**Figure 2.**
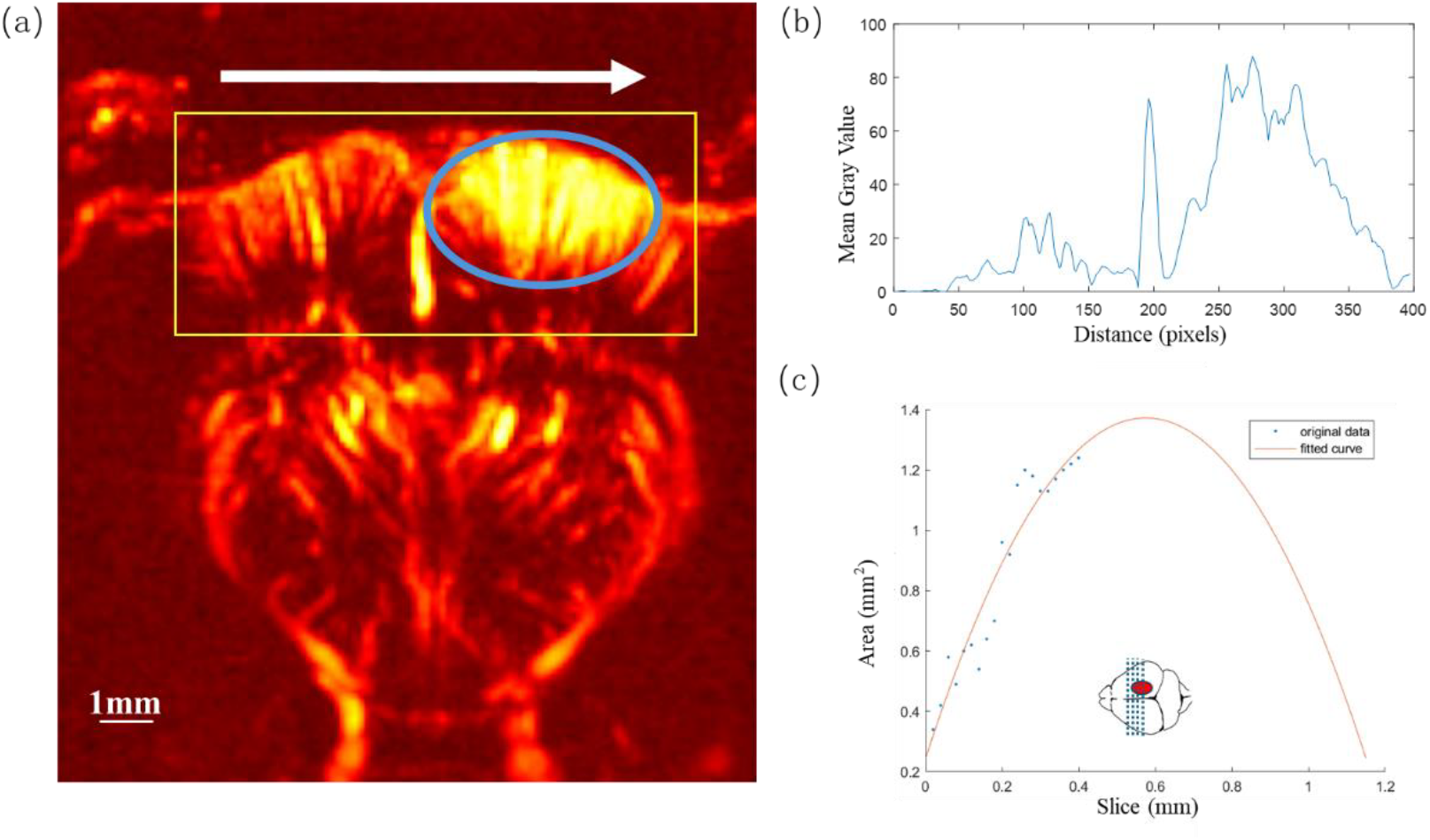
(a) fUS diagram of the coronal section of the rat brain, with a high-signal area indicated by the blue ellipse, suggesting increased blood flow velocity. (b) is a grayscale distribution of the yellow box in (a) along the direction of the white arrow. (c) Quantification of the area of the high-signal area in each section of the modeling area and proceed with curve fitting.

### 3. ULM evaluation of CSVD model in acute phase

Figs. 3(a)-(c) show the localization density of microbubbles in the coronal section of the rat brain in the acute phase of CSVD. The hollow arrow in Fig. 3(a) shows the modeling area on one day after modeling, and it can be seen that the number of long medullary arteries is less than that of the contralateral side, and the intervals between vasculars increase. When combined with Figs. 1(b), (d), and (e), it indicates that there are cerebral small vessel microinfarcts and microhemorrhage in the modeling area. On 3 days [Fig. 3(b)] and 6 days [Fig. 3(c)] after modeling, the number of long medullary arteries decreased compared with the number of long medullary arteries on one day after modeling, the intervals between vasculars increase gradually, the lengths become shorter, and the blood vessels become tortuous. In addition, the white matter area (blue double arrows) has widened.

**Figure 3.**
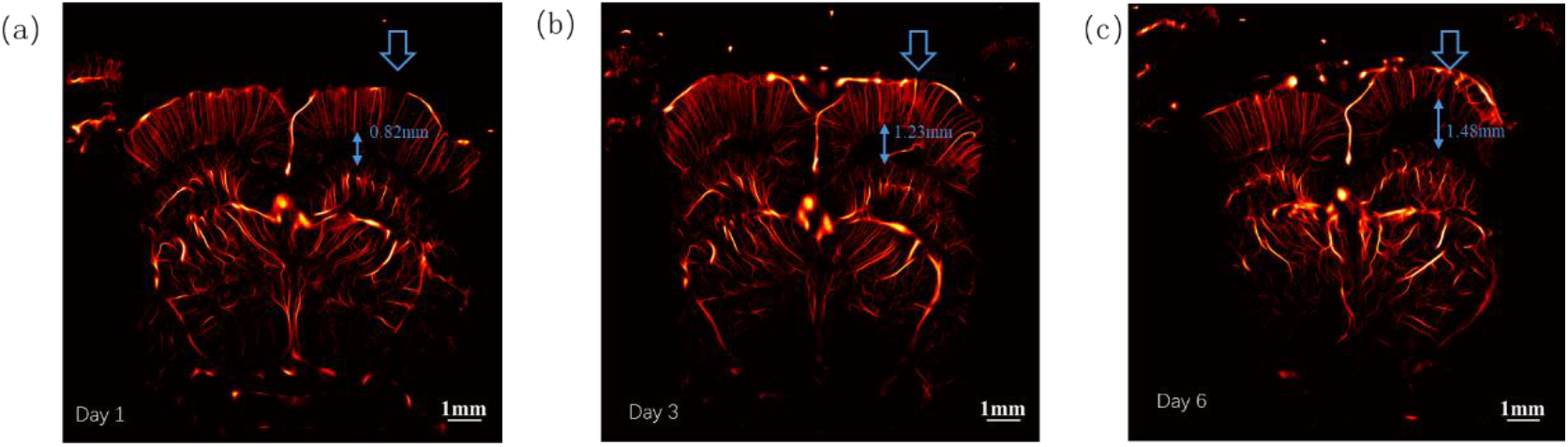
In the acute stage after modeling on (a) day 1, (b) day 3 and (c) day 6, the hollow arrows show the modeling area, and the number of long medullary arteries has reduced compared with the contralateral side. The intervals between vasculars have widened, the vessels are tortuous, and the width of the white matter area (blue double arrows) has widened.

### 4. Evaluation of micro blood flow velocity in the acute phase of the CSVD model

Fig. 4(a) shows the micro blood flow velocities (along depth direction) of CSVD in the acute phase of the coronal section of the rat brain, after 1, 3 and 6 days respectively. It is evident that the blood flow velocity in the modeled region has increased. Fig. 4(b) shows our quantification method. First, we draw two auxiliary lines parallel to meninges in the cortex on both sides of the brain. Then, two rectangles with the same size continuously move along the colored auxiliary lines on each side of the brain in the direction of white arrow. With the movements of the two rectangles, the average speeds of all pixels in the rectangles are calculated. Fig. 4(c) is line charts of the quantitative results of Fig. 4(b). The blue line shows the quantitative value on the left side (normal side) and the orange line shows the quantitative value on the right (modeling area involved). It can be seen that the blood flow speed on the normal side gradually decreases along arrow direction (from edge to middle), and the speed of the modeling side is lower than that of the normal side in the first half of auxiliary line, then higher than that of the normal side in the latter half of auxiliary line in the first three days, and completely higher than that of the normal side on the sixth day.

**Figure 4.**
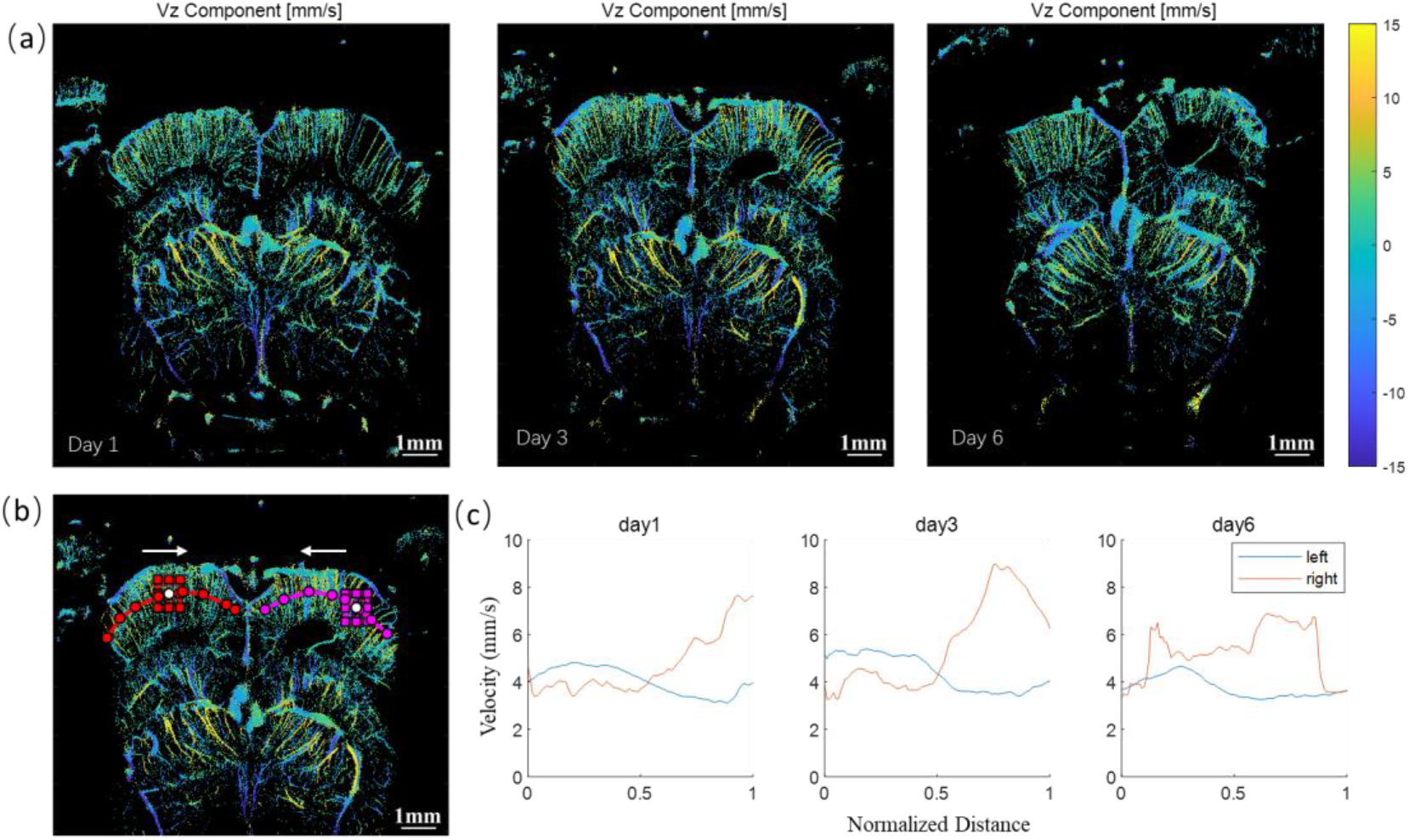
(a) Micro blood flow velocity diagrams (along z axis) of CSVD in the acute phase of the coronal section of the rat brain. (b) Two rectangles with a same size continuously move along the colored auxiliary lines on both sides of the brain in the direction of white arrow. (c) is the quantitative analysis of the cumulative average velocity in the two rectangles changing with their movements along the auxiliary lines in the direction of white arrow on (b).

## Discussion

In the early study^15^, we established rat CSVD model based on the pathogenesis of BBB dysfunction caused by vascular endothelial injury, using ultrasound combined with ultrasound contrast agent, which has a controllable degree of injury, controllable location, short modeling cycle and simple operation. At the same time, similar studies have emerged^17^. The experiments in this paper prove that the model can well simulate the image signs of STRIVE-2. The continuous observation of microvascular changes in the acute phase provides a piece of evidence on micro-circulation level for one of the pathogenesis of CSVD, namely based on BBB dysfunction caused by vascular endothelial injury.

BBB dysfunction is considered to be a core pathophysiological mechanism of CSVD^18,19^. As the disease progresses, it has formed four STRIVE-2 imaging features in our study: 1. recent subcortical infarct, 2. Cerebral microhemorrhage, 3. cerebral white matter hyperintensity (presumed vascular), 4. cerebral atrophy. Based on the BBB injury mechanism of CSVD, we induced BBB dysfunction through the combination of ultrasound and microbubbles and established a rat model consistent with the pathogenesis of CSVD. In our model, the reduction in the number of micro-vessels in the modeling area was seen on ultrasound ULM imaging [Fig. 1(c)], and microhemorrhage were seen in H&E staining [Figs. 1(d) and (e)]. On MRI T2 imaging [Fig. 1(b)], the modeling area showed mixed signals, which is consistent with small subcortical infarcts and microhemorrhage. White matter hyperintensities is the most common imaging feature of CSVD, which is characterized by the presence of multiple punctate, patchy, or mixed hyperintensities within the periventricular and subcortical white matter regions in hyperintense on T2-weighted imaging sequences. As shown by the hollow arrows in Fig. 1(a), the irradiated area is white matter hyperintensity, located around the lateral ventricles. CSVD is associated with cortical atrophy, as indicated by the solid arrow in Fig. 1(a), which highlights the thinning of the cerebral cortex in the region of interest.

Neuroimaging markers of CSVD, as delineated by the Strive-2 based MRI criteria, predominantly reflect the distal cerebral parenchymal sequelae of vascular pathology, rather than the vascular disease itself. CSVD is associated with a spectrum of clinical manifestations, including cognitive impairment. However, the relationships between CSVD-specific markers and clinical features such as cognition are inconsistent, suggesting that traditional markers may not fully encapsulate the nexus between small vessel lesions and significant outcomes. This inconsistency underscores the complexity of CSVD and the need for a more nuanced understanding of its pathophysiological mechanisms We have integrated traditional neuroimaging markers of CSVD with ultrasound multimodality to comprehensively assess intracranial small vessel hemodynamics, leveraging the distinct advantages of each modality to probe the vascular pathology more deeply. FUS employs high framerate multi-angle plane-wave imaging synthesis technology to achieve ultrafast imaging, which is capable of capturing the majority of blood flow signals within the brain with high sensitivity and spatial resolution. As depicted in Fig. 2(a), the coronal section of the rat brain exhibits an enhanced blood flow signal within the region of interest, demarcated by the blue circle. Fig. 2(b) illustrates the gray value distribution along the direction indicated by the white arrow within the yellow box. Furthermore, Fig. 2(c) presents the quantification of the molding area within the coronal section of the rat brain. The distinct advantage of fUS imaging lies in its rapid acquisition speed, which facilitates a preliminary estimation of the extent of brain pathology modeling. Moreover, fUS elucidates the increased microvascular blood flow velocity and the dysfunction of microvascular autoregulation during the acute phase of CSVD. The disadvantage of fUS is that it may not suffice to capture the intricate details of the microvasculature within the brain.

ULM is capable of capturing high-resolution blood flow images with a resolution finely up to 10 microns in live animals^20^. Fig. 3(a) is the localization density map of microbubbles in the coronal section of the rat brain one day after modeling, and the arrows show the modeling area, which shows that the number of long medullary arteries is lower than that of the contralateral side, the space between blood vessels widens. As illustrated in Figs. 3(b) and (c), at the 3-day and 6-day post-modeling, there was a significant decrease in the count of long medullary arteries. Additionally, the intervascular spaces were observed to dilate, the lengths of the arteries were notably reduced, and the course of the blood vessels exhibited pronounced tortuosity. Furthermore, the region of white matter, as indicated by the blue double arrows, exhibits an expansion. The cerebral microvasculature is supplied by two distinct arterial sources: the terminal branches of the leptomeningeal arteries within the subarachnoid space, and the perforating branches that arise directly from the major intracranial vessels^21^. The two arterial systems traverse in divergent directions, with one system penetrating the cerebral cortex and the other the deep medulla^22,23^. They ultimately converge within the subcortical white matter, establishing a critical watershed zone. In this region, the arterioles are exclusively terminal branches, lacking anastomotic connections, and exhibit a low vascular density^24,25^. Consequently, they are susceptible to localized circulatory ischemia and hypoperfusion. Acute ischemic events in these areas are associated with an increased risk of lacunar infarction or cerebral microhemorrhage, while chronic ischemia can result in white matter lesions.

ULM can not only obtain ultra-high-resolution blood flow images on live animals, but also obtain functional information through microvascular velocity analysis. Fig. 4(a) presents the velocity profile of CSVD in the acute phase, as observed in the coronal section of the rat brain. Figs. 4(b) and (c) depict the temporal changes in blood flow velocity: on the unaffected side, the velocity progressively diminished. In contrast, on the modeled side, the blood flow velocity was initially lower than that of the unaffected side during the initial three days, subsequently surpassing it, and by the sixth day, it was markedly higher than the velocity on the unaffected side. In general, the middle cerebral artery (MCA) possesses a greater diameter compared to the anterior cerebral artery (ACA)^26^. The MCA predominantly serves the lateral and deep structures of the cerebral hemisphere, whereas the ACA primarily supplies the medial regions of the frontal and parietal lobes. Consequently, the MCA’s blood supply territory is more extensive than that of the ACA. In terms of the energy consumption of brain functional areas, such as the analysis of the primary motor cortex, the cortical representation of muscle control exhibits an approximate inverted topographical arrangement, with the body’s motor map positioned in a ‘head-down’ configuration, although the facial representation remains ‘upright’^27^. The larger the primary motor cortex area is, the more precise control over muscles it provides, and the greater the corresponding oxygen consumption. Studies suggest that indicates that under normal circumstances, microvascular blood flow to the parietal lobe gradually decreases. After CSVD, the microvascular blood flow regulation dysfunction was impaired, and the modeling area showed abnormal perfusion.

This CSVD model, predicated on the disruption of the BBB due to vascular endothelial injury, effectively emulates the majority of neuroimaging signatures associated with CSVD. The application of fUS and ULM in the acute stage of CSVD has provided compelling evidence at the microvascular level, suggesting that BBB dysfunction is a central pathophysiological mechanism of CSVD. Then it provides a straightforward, efficient, and widely applicable tool and continuous monitoring method for the further investigation of CSVD from the macroscopic to the microscopic circulation level, with significant long-term implications for CSVD diagnosis and treatment.

## Statements and Declarations

This work was supported in part by the National Natural Science Foundation of China (Nos.82372561), Natural Science Foundation of Inner Mongolia Autonomous Region(Nos. 2022QN08013), Inner Mongolia Medical University university-level joint project plan(Nos. YKD2021LH071).

## Reference

1. Rincon F, Wright CB. Current pathophysiological concepts in cerebral small vessel disease. Front Aging Neurosci. 2014;6:24. doi:10.3389/fnagi.2014.00024.

2. Wong SM, Jansen JFA, Zhang CE, et al. Blood-brain barrier impairment and hypoperfusion are linked in cerebral small vessel disease. Neurology. 2019;92(15):e1669–e1677. doi:10.1212/WNL.0000000000007263.

3. Caplan LR. Lacunar infarction and small vessel disease: pathology and pathophysiology. J Stroke. 2015;17(1):2–6. doi:10.5853/jos.2015.17.1.2.

4. Nezu T, Hosomi N, Aoki S, et al. Endothelial dysfunction is associated with the severity of cerebral small vessel disease. Hypertens Res. 2015;38(4):291–297. doi:10.1038/hr.2014.162.

5. Hassan A, Hunt BJ, O’Sullivan M, et al. Homocysteine is a risk factor for cerebral small vessel disease, acting via endothelial dysfunction. Brain. 2004;127(Pt 1):212-219. doi:10.1093/brain/awh023.

6. Hainsworth AH, Oommen AT, Bridges LR. Endothelial cells and human cerebral small vessel disease. Brain Pathol. 2015;25(1):44–50. doi:10.1111/bpa.12231.

7. Van Agtmaal MJM, Houben AJHM, Pouwer F, et al. Association of microvascular dysfunction with late-life depression: a systematic review and meta-analysis. JAMA Psychiatry. 2017;74(7):729–739. doi:10.1001/jamapsychiatry.2017.0984.

8. Li Q, Yang Y, Reis C, et al. Cerebral small vessel disease. Cell Transplant. 2018;27(12):1711–1722. doi:10.1177/0963689718771213.

9. Wardlaw JM, Smith EE, Biesseels GJ, et al. Neuroimaging standards for research into small vessel disease and its contribution to ageing and neurodegeneration. Lancet Neurol. 2013;12(8):822–838. doi:10.1016/S1474-4422(13)70124-8.

10. Ye JY, Wang Z, Gong YT, et al. Neuroimaging standards for research into cerebral small vessel disease (STRIVE-2)—advances since 2013. Chin J Stroke. 2023;18(10):1160–1174.

11. Wardlaw JM, Smith C, Dichgans M. Mechanisms of sporadic cerebral small vessel disease: insights from neuroimaging. Lancet Neurol. 2013;12:483–497. doi:10.1016/S1474-4422(13)70060-0.

12. Wardlaw JM, Sandercock PA, Dennis MS, et al. Is breakdown of the blood-brain barrier responsible for lacunar stroke, leukoaraiosis, and dementia? Stroke. 2003;34:806–812. doi:10.1161/01.STR.0000063376.10803.76.

13. Hainsworth AH, Fisher MJ. A dysfunctional blood-brain barrier and cerebral small vessel disease. Neurology. 2017;88(5):420–421. doi:10.1212/WNL.0000000000003628.

14. Zhang CE, Wong SM, Van de Haar HJ, et al. Blood-brain barrier leakage is more widespread in patients with cerebral small vessel disease. Neurology. 2017;88(5):426–432. doi:10.1212/WNL.0000000000003627.

15. Ma H, Yang Y, Gao M, et al. A novel rat model of cerebral small vessel disease and evaluation by super-resolution ultrasound imaging. J Neurosci Methods. 2022;379:109673. doi:10.1016/j.jneumeth.2022.109673.

16. Hingot V, Brodin C, Lebrun F, et al. Early ultrafast ultrasound imaging of cerebral perfusion correlates with ischemic stroke outcomes and responses to treatment in mice. Theranostics. 2020;10(17):7480–7491. doi:10.7150/thno.44233.

17. He Y, Yang J, Hu F, et al. A new method for preparing a rat intracerebral hemorrhage model by combining focused ultrasound and microbubbles. Animal Model Exp Med. 2023;6(2):103–110. doi:10.1002/ame2.12303.

18. Wardlaw JM, Smith C, Dichgans M. Small vessel disease: mechanisms and clinical implications. Lancet Neurol. 2019;18:684–696. doi:10.1016/S1474-4422(19)30079-1.

19. Wardlaw JM. Blood-brain barrier and cerebral small vessel disease. J Neurol Sci. 2010;299:66–71. doi:10.1016/j.jns.2010.08.042.

20. Liu JB. Advances in deep learning-based ultrasound microscopy of microvasculature: basic and clinical research. Adv Ultrasound Diagn Ther. 2024;8(3):93.

21. Tariq N, Khatri R. Leptomeningeal collaterals in acute ischemic stroke. J Vasc Interv Neurol. 2008;1(4):91–95. doi:10.1016/j.jvir.2007.12.453.

22. Bogousslavsky J, Regli F. Centrum ovale infarcts: subcortical infarction in the superficial territory of the middle cerebral artery. Neurology. 1992;42:1992–1998. doi:10.1212/WNL.42.10.1992.

23. Read SJ, Pettigrew L, Schimmel L, et al. White matter medullary infarcts: acute subcortical infarction in the centrum ovale. Cerebrovasc Dis. 1998;8:289–295. doi:10.1159/000015866.

24. Gandolfo C, Del Sette M, Finocchi C, et al. Internal borderzone infarction in patients with ischemic stroke. Cerebrovasc Dis. 1998;8:255–258. doi:10.1159/000015864.

25. Lee PH, Bang OY, Oh SH, Joo IS, Huh K. Subcortical white matter infarcts: comparison of superficial perforating artery and internal border-zone infarcts using diffusion-weighted magnetic resonance imaging. Stroke. 2003;34(11):2630–2635. doi:10.1161/01.STR.0000097609.66185.05.

26. Gibo H, Carver CC, Rhoton AL Jr, et al. Microsurgical anatomy of the middle cerebral artery. Journal of Neurosurgery. 1981;54(2):151 – 169.

27. Schieber MH. Constraints on Somatotopic Organization in the Primary Motor Cortex. Journal of Neurophysiology. 2001;86(5):2125 – 2143. doi:10.1152/jn.2001.86.5.2125.

